# Loss of cholesterol in Junctional Epidermolysis Bullosa skin identifies a key role for Laminin-332 in actomyosin mediated cholesterol transport

**DOI:** 10.1101/2023.09.10.557030

**Authors:** Eleri. M. Jones, Emanuela Camera, Piotr Parzymies, Supatra.T. Marsh, Ryan.F. O’Shaughnessy, Monique Aumailley, John. A. McGrath, Edel.A. O’Toole, Matthew Caley

## Abstract

Individuals with Junctional Epidermolysis Bullosa (JEB), a rare genetic skin disease characterised by loss of function mutations in the Laminin332 (Lam332), do not survive beyond their first birthday. Here we report that loss of Lam332 leads to absence of cholesterol lipid from the epidermis *in vitro* and *in vivo*. Using 3D skin equivalents, a JEB mouse model and JEB patient samples we confirmed changes in epidermal lipid synthesis, which was further explored using lipidomics. Cholesterol biosynthesis genes were increased with loss of Laminin-332 in vitro, however a decrease in immunofluorescence lipid staining was observed. Cholesterol transport in Laminin-332 knockdown keratinocytes was revealed to be disrupted, which in keratinocytes is dependent on the actomyosin network. In conclusion these findings suggest a role for the basement membrane protein Laminin-332 in lipid metabolism in the skin, and a broader role for epidermal homeostasis and barrier formation. Restoration of cholesterol transport in epidermal keratinocytes of JEB patients offers the potential to improve their skin barrier.

## INTRODUCTION

Junctional Epidermolysis Bullosa (JEB) is a rare genetic skin disease characterised by skin fragility, failure to thrive and mechanically induced blistering^1^. Babies diagnosed with this form of EB generally do not survive beyond their first birthday^2^. Why this form of EB is so severe compared to recessive dystrophic epidermolysis bullosa (RDEB), despite the comparable depth of blisters and skin fragility, is poorly understood. The most severe form of JEB is caused by loss of function mutations in genes encoding the basement membrane protein Laminin332 (Lam332)^3^. Lam332 is a secreted extracellular glycoprotein composed of three genetically distinct chains α3, β3 and γ2 ^4,5^. It is an essential component of the dermal-epidermal junction, acting as a bridge between the hemidesmosomes attaching the basal keratinocyte to the basement membrane and collagen type VII in anchoring fibrils anchoring the basement membrane to the dermis.

The outermost layer of the epidermis, the stratum corneum, consists of differentiated keratinocytes in a neutral lipid rich matrix. This lipid matrix has an unique organisation consisting of ceramides, cholesterol and free fatty acids, which is important for the skin’s barrier function preventing water loss and entry of extrinsic pathogens ^6–8^.

In this study we developed a stable knockdown of each of the Lam332 chains (shLAMA3, shLAMB3 and shLAMC2) in an nTERT immortalised keratinocyte cell line to study any changes with loss of Lam332, as representative models of human JEB. Here we report that loss of Lam332 leads to lipid changes within the epidermis, specifically absence of cholesterol from the plasma membrane (PM) and epidermis in 3D in vitro models of JEB. The loss of skin cholesterol is due to a defect in cholesterol transport, which in keratinocytes we identify to be dependent on the actomyosin network, which is disrupted by loss of Lam332. In conclusion these findings suggest a role for Lam332 in lipid metabolism in the skin and a broader role in epidermal homeostasis and barrier formation. Restoration of cholesterol transport in JEB patients offers the potential to improve the skin barrier.

## RESULTS

### Loss of Laminin332 upregulates cholesterol biosynthesis genes

The α3 subunit of Lam332 was successfully knocked down with siRNA in primary keratinocytes (Supplementary Fig. 1a) with greater than 95% down-regulation compared to controls. Analysis by RNAseq, comparing these cells to siControl cells, generated a dataset of differentially expressed genes. Over representation analysis of this dataset (DAVID GO Terms, Supplementary Table. 2) identified a subset of genes involved in lipid biosynthesis that were significantly overrepresented in the siLamα3 gene list (Fig. 1a and b).

**Figure 1.**
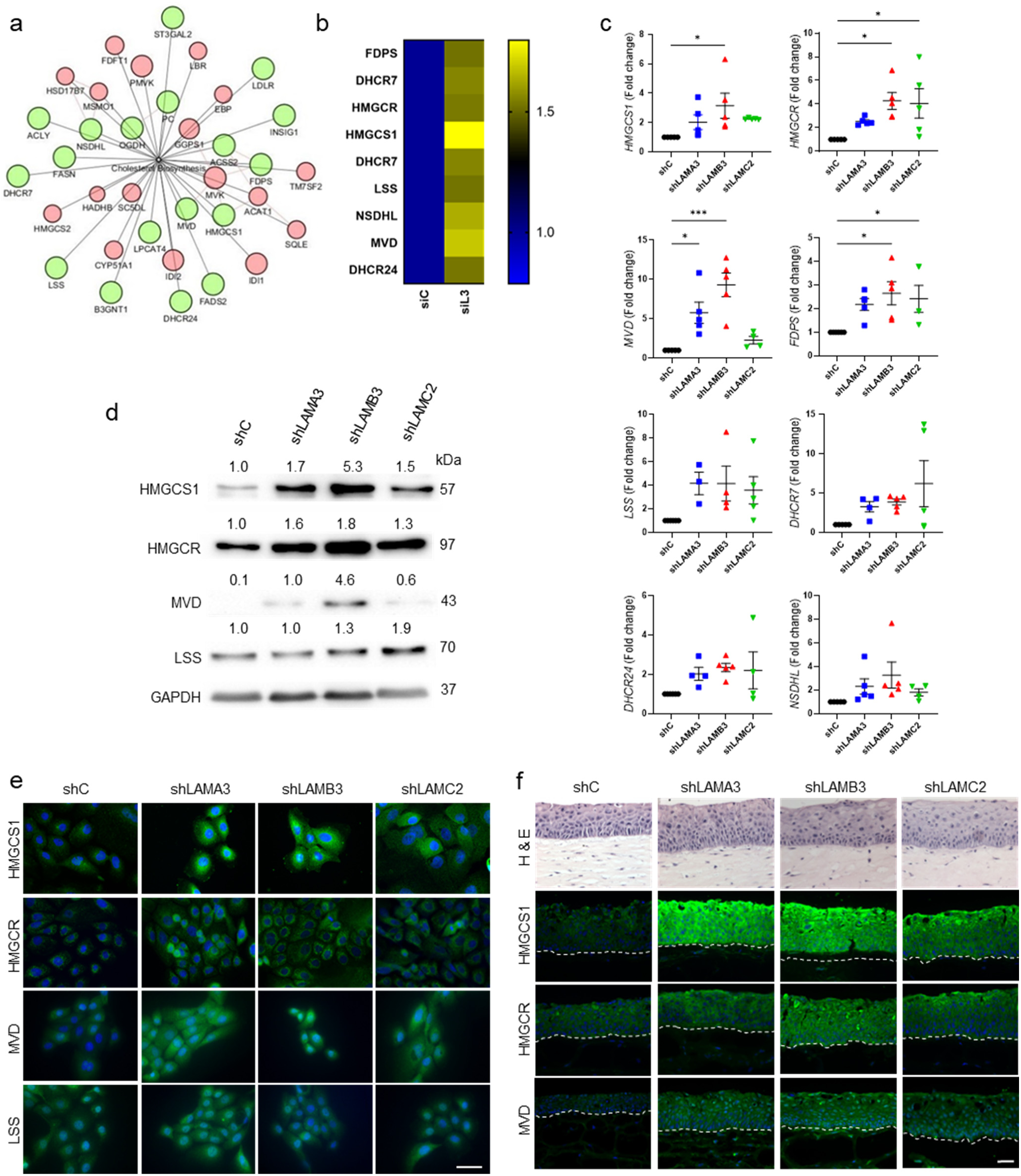
Loss of Laminin332 upregulates Cholesterol biosynthesis genes. **a)** DAVID clustering analysis of RNA sequencing data for siLamA3 gene knockdown. **b)** Fold change of cholesterol biosynthesis genes with loss of basement membrane protein Lam332. **c)** Real-time quantitative reverse transcription analysis of cholesterol biosynthesis genes in Laminin332 knockdown cells. Statistical analysis was performed using one-way ANOVA and compared to shC. *p<0.05, ***p<0.001. **d)** Western blotting analysis of HMGCS1, HMGCR, MVD and LSS in control (shC) and Lam332 knockdown (shLamA3, shLamB3, shLamC2) cells. GAPDH was used as an internal control of protein loading. For densitometry analysis, results were normalized to GAPDH and are expressed as fold induction over shC. **e)** Immunofluorescence staining of HMGCS1, HMGCR, MVD and LSS in cells (scale bar = 50µm). **f)** Haematoxylin & eosin and immunofluorescence staining of HMGCS1, HMGCR and MVD in 3D skin equivalents (scale bar = 50µm).

These observations were confirmed by generating stable knockdowns of each chain of Lam332 (shLAMA3, shLAMB3 and shLAMC2; Supplementary Fig. 2a) in an nTERT immortalised keratinocyte cell line. Upregulation of cholesterol biosynthesis pathway genes (*HMGCS1, HMGCR, MVD, LSS, FDPS, NSDHL, DHCR7* and *DHCR24*) was confirmed by real-time qPCR (Fig. 1c). The increase in *HMGCS1*, *HMGCR*, *MVD* and *LSS* was confirmed at protein level by western blotting (Fig. 1d) and immunofluorescence (Fig. 1e). 3D skin equivalent cultures were generated with shC, shLAMA3, shLAMB3, shLAMC2 and siC and siLamA3 keratinocytes and validated for loss of each Lam332 chain (Supplementary Fig. 2b and 1b). Cholesterol biosynthesis genes were upregulated in the 3D skin equivalent models simulating JEB (Fig. 1f and Supplementary. 1b).

An inducible mouse model of Laminin-α3^9^ (*Lama3*^flox/flox^/K14^CreERT^ mouse; KO) also showed changes in cholesterol biosynthesis genes. Immunostaining of skin sections from these mice revealed an upregulation of *Hmgcs1*, *Hmgcr* and *Mvd* compared to control (Ctrl) *Lama3*^wt/flox^/K14^CreERT^ mouse (Fig.2a). Expression of cholesterol biosynthesis genes was increased in both *LAMA3* (n=4) and *LAMB3* (n=3) mutant JEB patient samples compared to control skin (n=5) (Fig. 2b). Patient details are in Supplementary Table 3.

**Figure 2.**
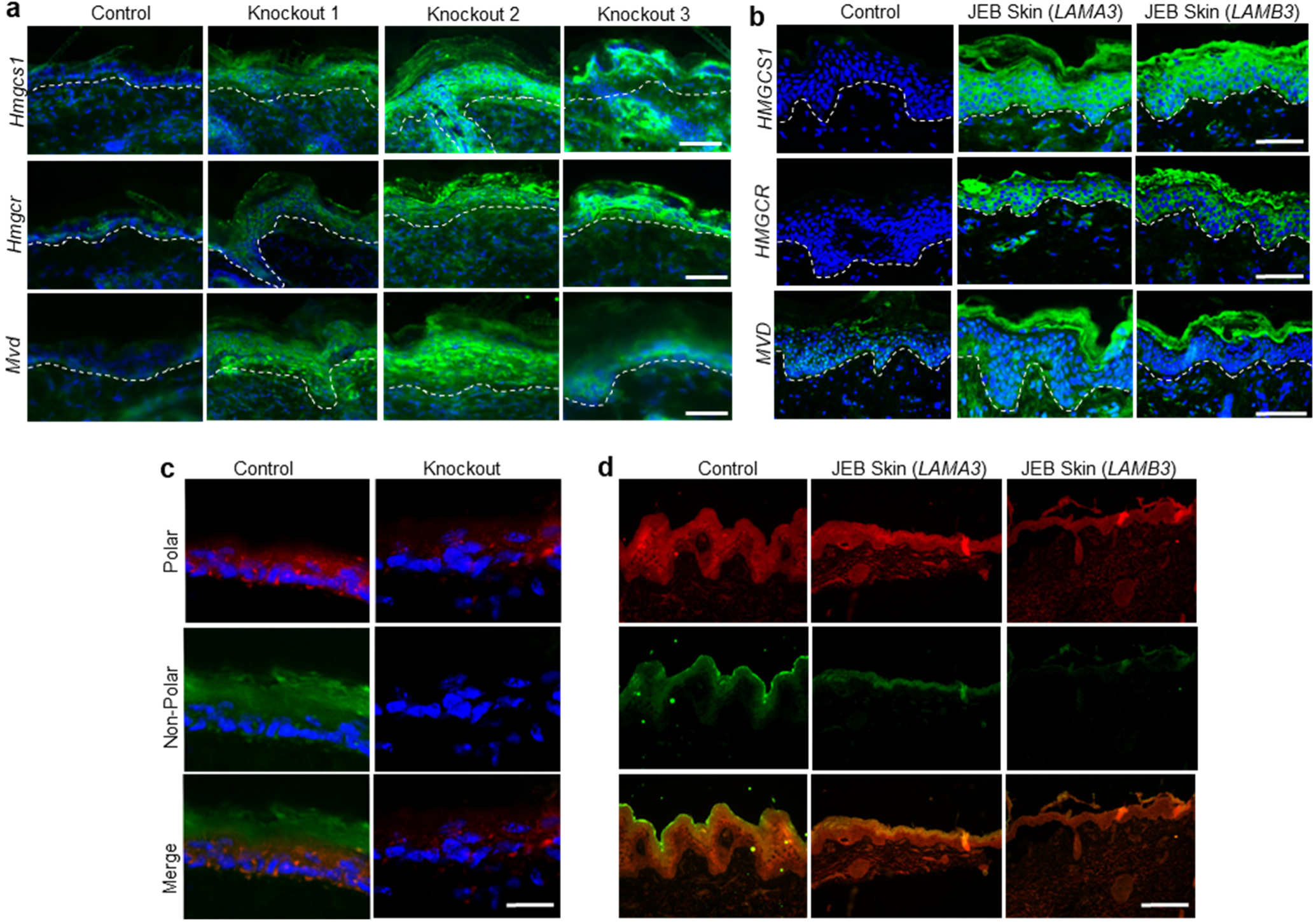
Cholesterol biosynthesis and lipid changes in *Lama3*^flox/flox^/K14-CreERT mouse and JEB patient skin. **a)** Immunofluorescence staining of cholesterol biosynthesis genes were performed in skin samples of a tamoxifen-treated heterozygous *Lama3*wt/flox/K14-CreERT (control) mouse and *Lama3*flox/flox/K14-CreERT (knockout). **b)** Representative images of control skin and JEB patient skin (with both *LAMA3* and *LAMB3* mutations stained for cholesterol biosynthesis. The lipid fluorescent stain Nile Red used to show both polar (red) and non-polar (green) lipids in **c)** mouse and **d)** JEB patient skin. Dotted line delineates the epidermis and dermis. DAPI (4’,6-diamidino-2-phenylindole) was used as a nuclear stain. Images shown are representative of 6 mice per group. JEB patient samples representative of 6 patients (3 with *LAMA3* mutation and 3 with *LAMB3* mutation). Scale bar = 50 µm

### Loss of Laminin332 results in changes in skin lipids

Nile red staining of siLamA3 skin equivalents (Supplementary Fig.1b), *Lama3* KO mouse (Fig. 2c) and JEB patient skin (Fig. 2d) revealed reduced polar and non-polar (barrier) lipids compared to respective controls. Loss of the basement membrane protein Collagen type VII, which results in another blistering disease RDEB ^10,11^ also demonstrated some lipid changes in 3D in vitro models (Supplementary Fig 3). A loss of barrier function was demonstrated in siLamA3 skin equivalents with failure to retain Lucifer yellow dye compared to siC (Supplementary Fig. 1b). To explore the epidermal lipid profile with loss of Lam332 we performed GC-MS in conjunction with untargeted/targeted LC-MS of *Lama3* mouse epidermis and Lam332 whole skin equivalents (epidermis and dermis). Untargeted LC-MS demonstrated substantial differences in comprehensive lipid profiles between control and *Lama3* KO mouse epidermal extracts in both positive (Fig. 3) and negative (Supplementary Fig. 4) ion mode (+/-ESI). Volcano plot analysis revealed significantly different lipid entities between samples with a fold change higher than 1.5 (Fig. 3b and Supplementary Fig 4b). Targeted analyses indicated a downregulation of glycerolipids, and a small percentage of sphingolipids and phospholipids in the *Lama3* KO compared to control mouse. In contrast, sphingolipids accounted for most upregulated lipids, followed by wax esters, glycerolipids, ethanolamines and phospholipids (Fig. 3c, d and Supplementary Fig. 4c). Cholesterol esters (CE) and cholesterol sulfate (CS) were quantitatively determined by LC-MS whereas cholesterol and desmosterol were analysed from the same lipid extracts by GC-MS. No significant changes were observed in cholesterol and desmosterol, but a significant upregulation of CS and downregulation of CE(18:1) was observed in the *Lama3* KO epidermis (Fig. 4e-h).

**Figure 3.**
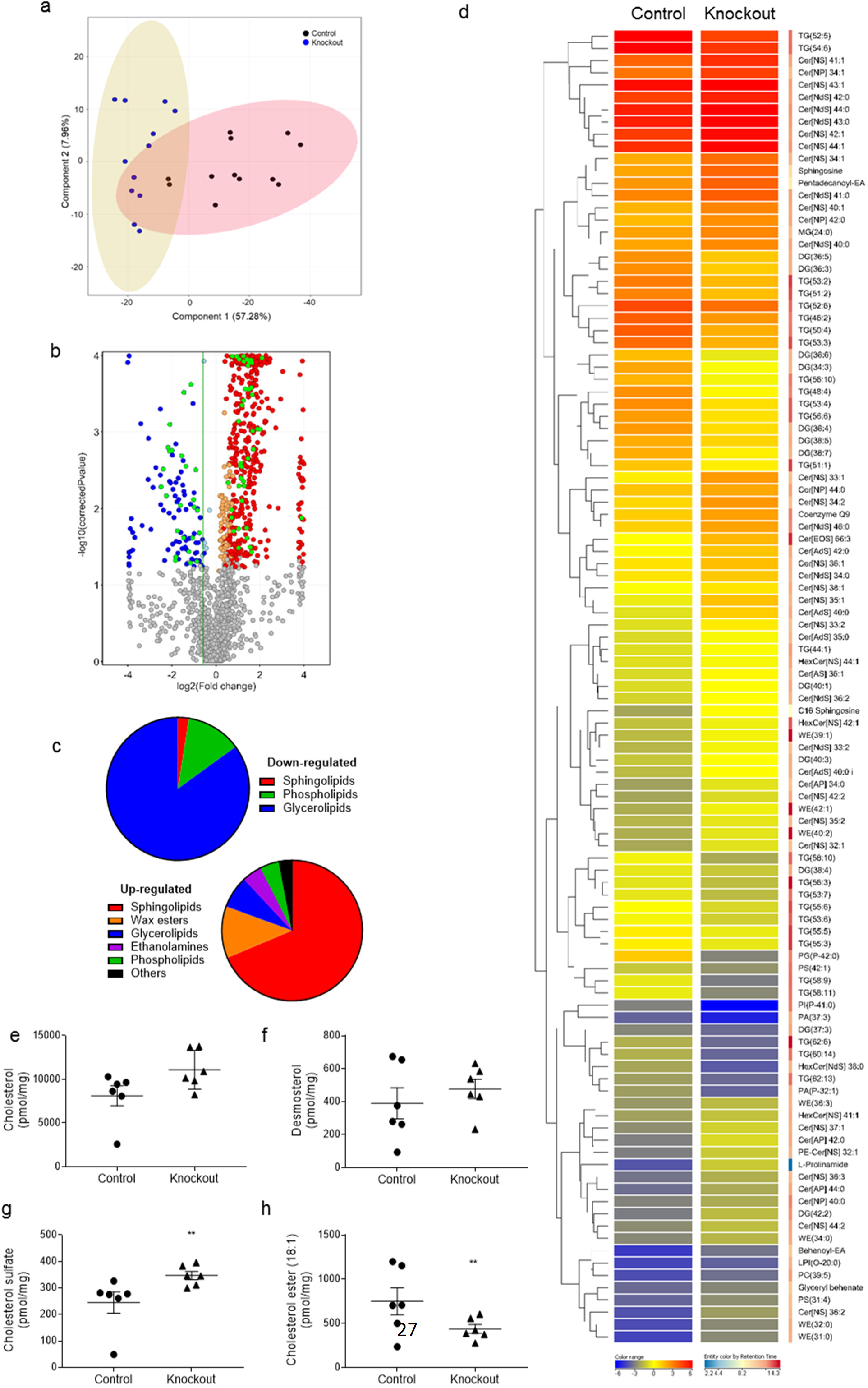
Lipidomic analysis of Lama3flox/flox/K14-CreERT mouse. Analysis from positive ion mode (+ESI). **a)** Principal component analysis (PCA) illustrating the variances between the two sample groups. Within the first two principal components 65.24% of the total variance was explained, with 57.28% in the first dimension and an additional 7.96% in the second dimension. **b)** Volcano plot analysis displaying significantly different regulated lipid species between Lama3 KO mice vs Control (Ctrl). The log2 of the fold change (FC) values were plotted on the x-axis, whereas the −log10 of the t-test p-values were plotted on the y-axis. The dots corresponding to each entity were coloured according to the FC, with red indicating significantly upregulated lipids and blue indicating significantly downregulated (based on p < 0.05 and FC > 1.5); green dots highlight the 107 lipid molecules further analysed in figures c and d. **c)** Further analysis revealed the main lipid sub-groups down-regulated (sphingolipids, phospholipids and glycerolipids) or upregulated (sphingolipids, wax esters, glycerolipids, ethanolamines, phospholipids and others) in KO vs Ctrl mouse groups and **d)** Integrated hierarchical clustering analysis of 107 lipid species through targeted MS/MS which were significantly different with a FC >1.5 between KO and Ctrl. Ceramide and triglyceride annotations were confirmed upon generation of MS/MS spectra. Colour scale (solid blue to solid red) reflects Log 2 values and colour scale (blue to red) shows retention time in the LC-MS analysis indicated in minutes. **e-h)** Quantitative analysis of cholesterol related lipid species cholesterol, desmosterol (with GC-MS), cholesterol sulfate and CE(18:1) (with LC-MS) respectively. Statistical analysis was performed using an unpaired t-test **p < 0.01 (n = 6).

**Figure 4.**
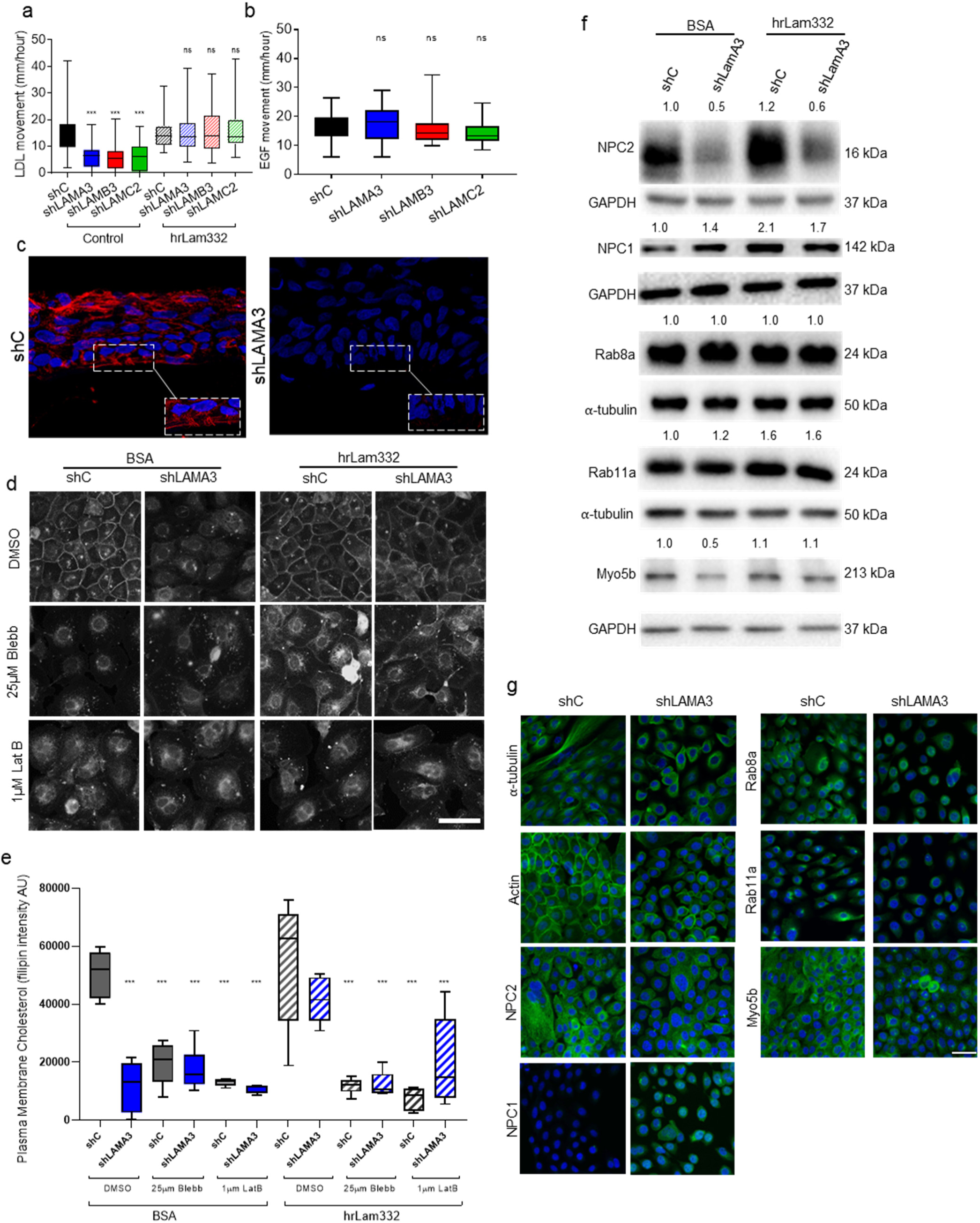
Cellular quantification and localisation of cholesterol with loss of Laminin332. **a)** Quantification of movement of fluorescently labelled dil-LDL (1,1′-dioctadecyl-3,3,3′,3′-tetramethylindo-carbocynanine perchlorate-LDL) in control and Lam332 knockdown cells cultured on BSA or hrLam332. **b)** Trafficking of fluorescent EGF as a control was carried out in both shC and Lam332 knockdown cells showing no significant difference. Videos (20 of each condition) were quantified using particle tracking on ICY software and statistical analysis performed. **c)** Representative images for actin cytoskeleton staining both in control and shLamA3 3D skin equivalents showing significant loss of actin fibres at the basement membrane in Lam332 skin equivalents. The inset in each panel represents a closer view of the basement membrane region d) Fluorescent filipin staining (which binds to free cholesterol within the cell) in control and shLamA3 knockdown keratinocytes cultured on BSA or hrLam332, with and without Blebbistatin and Latrunculin B. **e)** Images were quantified and results presented as intensity of plasma membrane filipin staining. **f)** Western blotting analysis of cholesterol transport pathway genes (NPC2, NPC1, Rab8a, Rab11a and Myo5b) in control and shLamA3 knockdown cells cultured on BSA or hrLam332. **g)** Immunofluorescence of α-tubulin, actin, NPC2, NPC1, Rab8a, Rab11a and Myo5b in control and shLamA3 knockdown cells. Statistical analysis for all above experiments was performed using one-way ANOVA, *p<0.05, **p<0.01 and ***p<0.001. Scale bar 50µm.

Similarly, analysis of Lam332 skin equivalents revealed considerable differences in the lipid profiles compared to control (Supplementary Fig 5a, b). Several species were universally modulated between the three laminin knockdown subtypes, 5 were commonly downmodulated and 23 were commonly upmodulated (Supplementary Fig.5c, d). However, each laminin chain knockdown resulted in variable lipid changes, specifically for the phospholipid sub-class (e.g. phosphatidylglycerol (PG), phosphatidylethanolamine (PE), phosphatidylinositol (PI)) (Supplemantary Fig. 5b, e). Quantitative analysis of cholesterol metabolites in skin equivalents revealed a significant upregulation of CS in the Lam332 knockdown samples (Supplementary Fig 5h).

### Loss of Laminin332 results in disrupted transport of cholesterol

Total cholesterol (TC) remained unchanged (Supplementary Fig. 6a) but CE was decreased (Supplementary Fig. 6b) with loss of Lam332 in cultured keratinocytes along with loss of cholesterol (nile red staining) in the epidermis (Fig. 2b), suggesting abnormal cholesterol transport in the skin. Uptake of LDL (low-density lipoprotein) cholesterol with loss of Lam332 was investigated by monitoring the binding and internalisation of 1,1′-dioctadecyl-3,3,3′,3′-tetramethylindo-carbocynanine perchlorate (dil-LDL) in Lam332 knockdown cells. There was no difference in LDL binding (Supplementary Fig. 6c & d) or LDL receptor expression in any of the cell lines (Supplementary Fig.6f). However, LDL uptake was significantly increased in Lam332 KD cells (Supplementary Fig.6c and e); suggesting a failure of the LDL to be recycled efficiently and therefore an accumulation within the cell. Furthermore, analysis of the LDL-receptor showed an accumulation of the precursor form (Supplementary Fig. 6f), suggesting an issue with the recycling of the mature receptor.

To further assess the transport of cholesterol within Lam332 knockdown cells, we imaged the movement of fluorescent dil-LDL and EGF by time lapse video microscopy over a 2-hour period. It showed reduced movement of dil-LDL in Lam332 knockdown cells compared to control (Supplementary Video 1-4), but no change in EGF movement (Supplementary Video 9-12). Particle tracking analysis on representative videos showed a significant reduction in LDL movement which was reversed when cells were cultured on human recombinant Lam332 (hrLam332) (Supplementary Video 5-8) (Fig. 4a). EGF movement, controlled by microtubules^12^, was unchanged in Lam332 knockdown cells (Fig. 4b), suggesting that cholesterol is transported within the cell through a different mechanism.

To further explore this defect in cholesterol transport, we investigated the proteins involved in the intracellular cholesterol transport pathway in shLAMA3 keratinocytes. Our analysis demonstrated that the levels of NPC2 and Myo5b were downregulated in shLAMA3 compared to shC (Fig. 4f and g), with a subsequent restoration when the cells were cultured on hrLam332. Interestingly, this restoration appeared to be more pronounced for Myo5b than NPC2 suggesting a potential time-dependent effect of the recombinant protein. Levels of NPC1, an NPC2-binding partner, were upregulated in shLAMA3 keratinocytes (Fig.4f) suggesting a potential compensatory mechanism. Myo5b partners, Rab8a and Rab11a (which regulate cholesterol recycling) were expressed at similar levels in shC and shLAMA3 keratinocytes (Fig.4f).

### Cholesterol movement is dependent on the actin cytoskeleton network

The actin cytoskeleton was analysed in Lam332 knockdown skin equivalents with phalloidin revealing a reduced intensity of filamentous actin throughout the epidermis, with a marked absence at the basement membrane, in all Lam332 knockdown skin equivalents (data shown for shLAMA3, Fig. 4c). Furthermore, filipin staining revealed a significant reduction in plasma membrane cholesterol in Lam332 knockdown cells compared to control, which was restored to control levels when cells were cultured on hrLam332 (Fig. 4d and e, Supplementary Fig.7). To verify whether cholesterol transport from the endoplasmic reticulum to the PM is dependent on the actin cytoskeleton we disrupted the actomyosin network, which revealed that cholesterol transport to the PM is dependent on the actin cytoskeleton as was visualised by the loss of PM cholesterol staining in control (shC) cells treated with blebbistatin and latrunculin B (Fig. 4d). This loss of cholesterol at the PM was not restored with hrLam332 (Fig. 4d).

Plasma membrane levels of the transmembrane β1 integrin in both of our cell lines grown on BSA and hrLam332 demonstrated a statistically significant decrease of integrin β1 in the shLAMA3 knockdown cells which was partially restored by the addition of hrLam332 (Supp Fig. 8a). The levels of both total and activated (y397) FAK were decreased in shLAMA3 keratinocytes with a complete restoration when the cells were grown on hrLam332 (Supp Fig. 8b).

## DISCUSSION

Laminin332 is an important component of the skin basement membrane, which is absent in JEB skin resulting in defective epidermal-dermal adhesion where patients suffer from severe blistering from birth^1^. Based on our observations and peer reviewed papers covering loss of lipids within the skin and their importance to the skin barrier^13–16^ we believe that the loss of lipids we have identified in JEB patient skin leads to an impaired skin barrier. Significant changes in epidermal lipids lead to an increase in trans epidermal water loss (TEWL) and barrier dysfunction in the epidermis^16^, reducing the skin’s ability to protect the body from external infections. We have demonstrated this impaired barrier *in vitro* and other groups have demonstrated that loss of Laminin 332 in mouse skin also impairs the skin barrier (through increased TEWL)^17^. Depletion of stratum corneum lipids is already associated with numerous skin diseases including atopic dermatitis (AD)^18^, ichthyosis^19^, xerosis^20^, acne vulgaris^21,22^ and aged dry skin^23^. Loss of Collagen VII (reduced or mutated in RDEB) also showed a decrease in epidermal lipids (Supplementary Fig 3), therefore studying other basement membrane proteins such as Collagen IV, VII or XVII, could reveal a lipid biosynthesis and metabolism defect in all EB diseases.

Lipidomic analysis revealed significant changes in epidermal lipids in both mouse and 3D skin equivalents with loss of Lam332. For mouse skin only the epidermis was analysed, whereas the skin equivalents were analysed whole, which included the HDF layer. The HDFs, however, were from the same donor in each shLAM332 skin equivalent and therefore unlikely to influence the lipidomic analysis presented here. The crosstalk and signal transduction between the epidermal and dermal cells may have a role within lipid metabolism and therefore further analysis of the fibroblasts from the *Lama3* KO mouse could reveal further insights into the lipid changes observed. Inflammation and immune responses are also regulated by lipids^24^ and in the mouse model could also influence lipid changes, these changes cannot be investigated in our current 3D skin equivalents. Although both disease models resulted in distinct lipid changes within the epidermis, both offer an insight into the role of Lam332 in formation of the lipid barrier. The *Lama3* KO mouse revealed a significant upregulation in sphingolipids. Ceramides are considered key in epidermal barrier maintenance^25,26^ and in regulating keratinocyte proliferation, differentiation and apoptosis^27^. Ceramides have been extensively studied in psoriasis, AD and ichthyoses^26^. An increase in specifically short chain ceramides was observed which has previously been shown to limit formation of an efficient lipid barrier in AD^28^. In the skin equivalent model however, it was changes in phospholipids that were highlighted with loss of Lam332. Phospholipids are essential for cellular dynamics^29^ and alterations may affect the epidermal barrier^30^. Significant changes in phospholipids have previously been associated with AD^31^. Evidence has shown that an increase in ceramides regulates cholesterol metabolism and reduces its translocation,^32^ and can also alter membrane dynamics reorganising the lipid domain resulting in actin cytoskeleton remodelling^33,34^. This imbalance of membrane lipids could have a significant effect on the defensive role lipids play in the epidermal barrier of the skin^35^.

Cholesterol is an important regulator of lipid organisation, and a complex cellular mechanism is required to maintain cholesterol levels in membranes^36^. A reduction in CE led to the discovery that cholesterol transport with loss of Lam332 was significantly impaired. Despite extensive research into Niemann-Pick disease, the most common cholesterol transport disorder^37^, there remains unresolved questions on how cholesterol moves within cells and how it is exported. In inflammatory skin diseases such as psoriasis, evidence has shown a role for cholesterol in IL17A signalling^38^. Furthermore, evidence suggests that lipid metabolism and cholesterol trafficking directly regulate the inflammatory pathways in macrophages^39^; suggesting that cholesterol could have multiple roles within the skin providing an adequate lipid barrier as well as maintaining the inflammatory response.

We describe for the first time that loss of Lam332 at the basement membrane directly results in defective transport of cholesterol to the epidermis due to instability of the actomyosin network. While intracellular cholesterol transport towards the plasma membrane has been widely studied in other cell lines including; squamous cell carcinoma (A431)^40^ rat hepatoma (CRL-1601) cell line^41^, Chinese hamster ovary cells^42^, fibroblasts^43,44^ including human skin fibroblasts^45^ the molecular mechanisms in skin keratinocytes are poorly understood^46^. We show that in keratinocytes, cholesterol movement is dependent on vesicle transport driven by the actin cytoskeleton. We further delved into the cholesterol transport pathway, briefly, extracellular cholesterol (carried by LDL) is internalised by the cell in the process of receptor-mediated endocytosis and subsequently de-esterified in the early endosome. Free cholesterol is transferred by the soluble NPC2 protein to the endosome membrane bound NPC1. The cargo is then delivered back to the plasma membrane by the activity of Rab8a or Rab11a GTPases and the actin bound Myo5b^40,47,48^. Myo5b has a well-known role in the final stages of cholesterol recycling^40^, whereby its deletion in a 3D human skin model perturbed the epidermal barrier function leading to a decrease in epidermal lipids and cell-cell junction proteins^48^. Furthermore, Myo5b KO mice show extensive wrinkling, characteristic of a defective epidermal barrier^49^. We did not observe downregulation of Rab8a or Rab11a in our shLAMA3 keratinocytes suggesting that the cell’s ability to produce lipid-transporting vesicles such as lamellar bodies is not compromised and that our phenotype is likely due to the cell’s inability to transport those vesicles efficiently.

Cholesterol recycled by actin dependent Myo5b has been shown to be preferentially delivered to the sites of focal adhesions^40^. Focal adhesion kinase (FAK) is a key component of focal adhesions playing a crucial role in the stabilisation of the actin cytoskeleton^50^. Our results demonstrated that shLAMA3 keratinocytes had lower levels of FAK (both total and activated, Supp. Fig 8), as well as lower levels of plasma membrane integrin β1, which is known to act as a Lam332 binding partner and is an important component of focal adhesions^51^. JEB patient keratinocytes show aberrant formation of actin containing focal adhesions^52^, and an abnormal distribution of integrin α6β4 along the basement membrane of JEB patient skin^53,54^. Lam332 is an important adhesion molecule, which along with integrin α6β4 and α3β1, BPAG2, and plectin^55^, contribute to anchorage of keratinocytes and formation of the hemidesmosome. Disruption of any of the Lam332 genes result in destabilisation of the hemidesmosome and importantly the cytoskeletal networks^56^. These results corroborate previous findings that loss of Lam332 results in a disrupted actin cytoskeleton^57^.

There is no cure for patients with JEB, while current treatment is aimed only at palliative care^58^. An additional disease burden to JEB patient’s quality of life has been identified as itch, with 100% of patients reporting issues with itch^59^. In other skin diseases associated with extreme itch the use of lipid containing emollients to temporarily restore the skin barrier has been shown to relieve itch^60,61^. While in other skin models the loss of skin lipids leads to an impaired barrier^62,63^, impaired wound healing^64^ and reduced microbial defense^65^. Improving any of these co-morbidities would improve the health of JEB patients. Targeting the lipid defect presented here, while it would not prevent the severe blistering or reverse skin fragility associated with JEB, it does offer the prospect of alleviating symptoms related to pain and itch by improving the severe barrier defect, whilst also offering the prospect of reducing the occurrent skin infections observed in JEB patients.

## MATERIALS AND METHODS

### Cell Culture

The human keratinocyte telomerase reverse-transcriptase (h/TERT)-immortalised N/TERT-1 cell line derived from clinically normal foreskin tissue and supplied by Professor James Rheinwald (Department of Dermatology, Harvard University Medical School, Boston, USA) was cultured in DMEM:Ham’s F12 (3:1) supplemented with 10% FCS, 1% L-glutamine (200 mM) and RM Plus (0.4 µg/ml hydrocortisone, 5 µg/ml insulin, 10 ng/ml EGF, 5 µg/ml transferrin, 8.4 ng/ml cholera toxin, and 13 ng/ml liothyronine). Human primary fibroblasts, isolated from fresh redundant foreskin, (HDF) were cultured in DMEM supplemented with 10% FCS and 1% L-glutamine (200 mM). All cells were cultured at 37°C and 5% CO_2_.

### Generation of shRNA-Transduced Cell Lines

For stable knock-down of Laminin332 chains, α3, β3 and γ2 in keratinocytes, the nTERT keratinocyte cell line was transduced with SMARTvector™ lentiviral particles (Thermo Fisher Scientific, Paisley, UK) of three different shRNA clones (Supplementary Table S3) targeting either laminin chains as well as non-targeting particles (negative control). N-TERT keratinocyte cells, seeded at 50% confluency, were incubated overnight with viral particles at a MOI (multiplicity of infection) of 2 diluted in complete DMEM:Ham’s F12 media supplemented with polybrene (5 µg/ml). Transduced cells were selected in puromycin (2.5 µg/ml) 48 hours after.

### siRNA Transfection

For Laminin-α3 chain knockdown, the nTERT keratinocyte cell line was transfected with a SMARTpool of four synthetic siRNAs (Dharmacon, UK), targeting the α3 chain or non-targeting siRNAs (siC) as a negative control (Supplementary Table 1). Cells were plated at 60% confluency and transfected with 4µg of Dharmafect1 (Thermo Fisher Scientific) transfection reagent and 12.5nM final concentration of each siRNA. Cells were isolated 48hrs post-transfection and protein isolated for western blot analysis or RNA extracted for sequencing analysis. siRNA transfected cells for epidermal models were also used after 48hrs post transfection and models were cultured as described below.

### *In vitro* epidermal models

*In vitro* 3D skin equivalents were prepared by adding 340µl of a Collagen/Matrigel containing 3.4×104 HDFs (166.6µl type I Collagen (Corning), 71.4µl of Matrigel (BD Biosciences), 34µl of 10x MEM, 34µl of FCS and 34µl of HDFs, resuspended at 1×106 in fibroblasts media) into a 12-well plate transwell insert (0.4µm pore size, Greiner). Gels were incubated for 1 hour to equilibrate and then keratinocytes (shC, shLAMA3, shLAMB3 or shLAMC2) were seeded on top at a density of 3.4×105 per gel. After 24 hours, gels were raised to the air-liquid interface and the models grown at an air/liquid interface for 14 days with daily media changes. L-ascorbic acid (Sigma) was added to the media at a concentration of 50µg/mL from day 8. The gels were harvested at day 14, either fixed in 4% paraformaldehyde (PFA) and embedded in paraffin or embedded in optimal cutting temperature (OCT) compound and frozen.

### Protein Analysis

For cell lysate immunoblot analysis, keratinocytes were lysed in 1x RIPA buffer (Fisher Scientific) supplemented with a protease-inhibitor-cocktail (Roche, UK). Lysates were subjected to 10% SDS-PAGE. The presence of protein was detected by immunoblotting using a primary antibody at 4°C overnight, and a horseradish peroxidase-coupled anti-mouse or - rabbit secondary antibody for 1 hour at room temperature and developed using Enhanced Chemi Luminescence. For densitometry analysis, the image analysis program Image J was used.

### qRTPCR

RNA extraction was performed using the RNeasy mini kit (Qiagen, Manchester, UK) according to the manufacturer’s instructions. cDNA was generated using the SuperScript® VILO™ cDNA Synthesis kit (Invitrogen, Paisley, UK). qPCR was carried out using KAPA SYBR® Universal qPCR Mastermix (Anachem, Luton, UK) on a StepOnePlus Real-Time PCR System (Thermo Fisher Scientific). Primer sequences are shown in Supplementary Table 5.

### Cholesterol quantification

Total cholesterol (TC) and free cholesterol (FC) were determined using the Cholesterol/Cholesteryl Ester Quantitation Assay (Abcam plc., Cambridge, UK), according to the manufacturer’s protocol. Briefly, samples (1 × 106 cells) were extracted with 200 µl of chloroform:isopropanol:NP-40 (7:11:0.1) using a microhomogenizer. After centrifugation at 13,000g for 10 min to remove insoluble material, the organic phase of samples was transferred to a new tube and dried in a vacuum for 30 min to remove chloroform. Dried lipids were then dissolved with 200 µl of the Cholesterol Assay Buffer. Fifty microlitres of samples and standards (1–5 ng) were added to 200µl of reaction mixture after 60 min incubation at 37°C the OD was measured at 570nm by an ELISA-reader. Cholesterol esters (CE) were determined by subtracting the value of FC from the TC.

### Filipin Fluorescence Staining

A cell-based Cholesterol Assay Kit from Abcam (Cambridge, UK) was performed in keratinocyte cells to visualize cholesterol by using Filipin III as a fluorescence probe of cholesterol. Briefly, 20,000 cells were seeded on glass coverslips in a 24 well plate and cultured in DMEM/F12 for 48 hours. After removal of culture medium from wells, cells were washed with PBS and fixed with PFA for 10 min and washed again with PBS (3 x 5 min). Filipin III solution was added to each well assayed and maintained in the dark for 45 min at room temperature. After washing (2 x 5 min PBS) fluorescence images were obtained by excitation at 350nm using a 40x oil objective.

### Fluorescence and Immunostaining

The use of archival human tissue sections was conducted according to Declaration of Helsinki principles and approved by the City and East NRES Committee approval number 05/Q0603/9. Skin sections were obtained of frozen human JEB and normal skin. Five μm-thick cryosections from skin equivalent cultures were cut in a cryostat, air-dried and fixed in 4% paraformaldehyde for 10 minutes before staining. Five μm-thick paraffin sections were de-paraffinized using xylene, hydrated in descending grades of ethanol to distilled water, and antigen retrieval was performed by heating samples in boiling 10 mM citrate buffer, pH 6.0, for 10 minutes.

Sections from skin equivalent cultures or patient skin were blocked at room temperature by incubating in blocking buffer (1% BSA (w/v) 2% FCS (v/v)) for 1 hour. Incubation with the primary antibody (see Supplementary Table 6) diluted in blocking buffer was performed at 4°C overnight. The secondary Alexa Fluor 568-red or 488-green, goat anti-rabbit or goat anti-mouse antibodies were added at a 1:250 dilution for 1 hour at room temperature. DAPI (300nM) was used as a nuclear stain. Images were photographed using a Leica epifluorescence microscope. Images were analysed using ImageJ and CellProfiler.

### Nile Red Lipid stain

For Nile Red lipid staining a stock solution containing 0.05% Nile Red (Sigma-Aldrich) in acetone was diluted to 2.5ug/ml with 75:25 glycerol:water, with Dapi (1:1000) followed by rapid vortexing. 20ul of solution was applied to air-dried cryosections and immediately covered with a coverslip. Green (488nm) and red (568nm) channels images were taken using an Epifluorescence Leica microscope.

### Permeability assay (Lucifer Yellow)

The 3D skin equivalents were incubated for 4 hours with 1 mM Lucifer Yellow (Sigma-Aldrich) solution on their upper surface, then PBS washed and cryo-embedded. Images were captured using an Epifluorescence Leica microscope with wavelength 488nm.

### Dil-LDL binding and uptake

Cells were incubated with 5µg/ml of dil-LDL for 30 minutes at 4°C (binding) or 30 minutes at 37°C (uptake) in epilife medium prior to fixation with 4% PFA and staining with DAPI for detection of nuclei. Images were taken on the Incell 2200 microscope and dil-LDL intensity was quantified. The total intensity for dil-LDL was divided by the number of nuclei per image to obtain an intensity per cell value.

Cells were seeded (3000 per well) in a 96 well plate, either coated with BSA or recombinant Laminin332 (BioLamina LN332-0502, diluted in Dulbecco’s Phosphate Buffered Saline, Sigma, D8662) and left to adhere overnight. Cells were starved of cholesterol for 24hrs in epilife culture media. Fluorescent dil-LDL was added at dilution of 1:200 for 10mins, cells were washed repeatedly with PBS-heparin (to remove any surface bound LDL). They were then incubated with DRAQ5 (nuclear dye) and ER-tracker in epilife media for live imaging. Cells were imaged at 40x every 4 minutes for 2 hours using the InCell 2200 and videos created with the software. LDL movement was tracked using ICY software and an average of 20 images analysed.

### Actin disruption

Cells seeded onto coverslips coated with either BSA or human recombinant Laminin332 were treated with blebbistatin (25µm) or latrunculinB (1µm) for 24 hours before being fixed and stained with filipin to visualise plasma membrane cholesterol.

### Flow cytometry

Cells were seeded at 200,000 cells in a T25 flask coated with 3 ml of 5µg/mL of BSA or hrLam332 (as previous) and incubated for 48 hours. Cells were trypsinised and 50,000 cells were placed in each FACS tube, centrifuged for 5min at 380g at 4°C. The supernatant was discarded, and the cells were resuspended in PBS containing APC mouse anti-human integrin-β1 antibody or the IgG1 K isotype control (1:100 dilution) and incubated for 15 minutes. 500µl of PBS was added to each tube, the cells were centrifuged in the same conditions and the supernatant was discarded. The cells were then resuspended in 300µl of DAPI dissolved in PBS (1:2,000) and analysed immediately using the BD FACS Canto II Flow Cytometry system. The geometric mean of plasma membrane integrin β1 intensity per single cell was then calculated using the FlowJo software and corrected for non-specific background signal using the isotype controls and DAPI only samples as negative controls.

### Sample processing for the analysis of epidermal lipids

For mouse samples (control n=6 and *Lama3* KO n=6), the epidermis was separated from the dermis by placing whole skin sections in dispase solution overnight at 4°C, while skin equivalents (shC n=3, shLAMA3 n=3, shLAMB3 n=3 and shLAMC2 n=3) were treated whole. Mouse epidermis or skin equivalents were weighed and lipids extracted with a chloroform/methanol mixture 2:1 after addition of the internal standard mixture containing SPLASH Lipidomix®,-LM6002 and d31Cer[NS] 34:1 (Avanti Polar Lipids, USA), and in-house mixed deuterated standards supplied by C/D/N isotopes and Toronto Research Chemicals, both from Canada (see Supplementary Table 7). Aliquots of dissolved lipid extracts were analysed by GC-MS for the quantification of cholesterol, desmosterol and free fatty acids. The dissolved lipid extracts were further analysed by untargeted LC-MS in positive and negative ion mode. The results of the untargeted approach were normalised by the internal standard d31Cer[NS] 34:1 and the weight of each sample. Quantitative results from both GC-MS and LC-MS were normalized by the mg of tissue weight and reported as pmol/mg.

### Gas chromatography-mass spectrometry

Gas chromatography coupled to electron ionization mass spectrometry (GC-MS) dual scan-selected ion monitoring was employed to determine quantitatively target compounds in the lipid extracts. Samples were analyzed with a GC 7890A coupled to the MS 5975 VL analyzer (Agilent Technologies, CA, USA). Quantitative analysis of sterols was performed as previously reported (Singh K, 2018). Briefly, dissolved lipid extracts were dried under nitrogen and derivatized with 50 µL BSTFA added with 1% trimethylchlorosilane (TCMS) in pyridine. To generate the trimethylsilyl (TMS) derivatives of sterols, the reaction was carried out at 60 °C for 60 minutes. GC separation was performed with the 30 m–0.250 (i.d.) GC DB-5MS UI fused silica column (Agilent Technologies, CA, USA), chemically bonded with a 5% diphenyl 95% dimethylpolysiloxane cross-linked stationary phase (0.25 mm film thickness). Helium was used as the carrier gas. Samples were acquired in scan mode by means of electron impact (EI) MS. Cholesterol and desmosterol were determined against d7cholesterol and d6desmosterol, respectively, with the MassHunter quantitative software (Agilent Technologies, CA, USA).

### Liquid chromatography-mass spectrometry

The chromatographic apparatus consisted of the 1260 Infinity II series LC system (Agilent Technologies, CA, USA). The stationary phase of the high-resolution reversed phase LC was a Zorbax SB-C8 Zorbax SB-C8 rapid resolution HT 2.1 × 100 mm 1.8 µm p.s. with a maximal operational backpressure at 600 Bar (Agilent Technologies, CA, USA). Lipid mixtures were eluted with a gradient of (A) 5 mM ammonium formate in water, (B) methanol, (C) acetonitrile, (D) isopropanol. The mobile phases were filtered through 0.45 µm glass filters and continuously degassed under vacuum. The elution program was as follows: A/B/C/D 60/28/8/40 at time 0 and held for 1 min, brought to A/B/C/D 1/70/20/9 in 10 min and held up to 20 min. The flow rate was maintained at 400 µL/min during the entire LC run. The column was thermostated at 60 °C. The injection volume was 0.20 µL. The injector needle was washed with the mobile phase in the wash port during the LC runs. The eluent outlet was connected to two different MS analyzers for the detection and characterization.

Accurate mass measurements in full MS and auto MS/MS were conducted with a G6545B series hyphenated QTOF (Agilent Technologies, USA) equipped with a JetStream Technology electrospray interface (ESI) operating in both positive and negative ion mode. Analytes eluted from the LC system were introduced into the Q-TOF apparatus at the operating chromatographic flow rate (see chromatographic conditions). Nitrogen was used as the nebulizing and desolvation gas. The temperature and the flow of the drying gas temperature were 200 °C, and 12 L/min, respectively. The temperature and the flow of the sheath gas were 350°C and 12 L/min, respectively. The nebulizer pressure was 40 psi. The capillary and the fragmentor voltage were 4000 and 180 V, respectively. Full scan mass spectra were acquired in the positive and negative ion modes in the range from m/z 100 to m/z 1600. To enhance accurate mass measurement for the ion species a reference solution of two compounds with m/z 121.050873 and 922.009798, in positive ion mode and m/z 112.995587 and 966.000725 in negative ion mode, was vaporized in continuum in the spray chamber by means of a separate nebulizer.

### Extraction of MS features

Molecular features, defined by an m/z, ration, retention time (RT) and signal intensity value, were extracted in profile mode within the 10 ppm mass window from the raw LC-MS data files using the untargeted or the targeted batch recursive feature extraction in the MassHunter Profinder software (Agilent Technologies, USA). Procedures and details can be found in ‘MassHunter Profinder Software Quick Start Guide. G3835-90027_Profinder_QuickStart’ on the Agilent Technologies webpage. The features extracted were exported into a compound exchange format (CEF) reporting RT, the accurate mass and the absolute abundance for each entity to be processed in the subsequent chemometric analysis as previously reported ^66,67^.

## Data analysis

Agilent Mass Profiler Professional (MPP version 15.1) was used to process the LC-MS untargeted and targeted data. RT were aligned by setting a RT window of 0.6 minutes, whereas m/z binning was performed by setting windows at 10 ppm. Absolute abundance of each entity was normalized by the absolute abundance of the d31Cer[NS] 34:1 internal standard. Data were filtered by frequency of detection, which reflects the number of samples that presented particular features. A frequency filter was applied to data extracted from MPP and only entities present in 100% of samples belonging to at least one of the investigated groups were retained for the statistical analysis. Fold changes of filtered entities were compared between groups volcano plots in the MPP tools. Fold changes with p values < 0.05 after the Bonferroni’s correction were considered as significant. Identification of entities within the MPP workflow was performed based on the METLIN Metabolomics Database (http://metlin.scripps.edu/) and the Lipid Annotator software (Agilent Technologies, CA, USA). Quantitative assessments of cholesterol sulfate and CE were performed with the labelled internal standards d7CHS, and d7CE (18:1), respectively.

## Supplementary Materials

Figure S1. Knockdown of Laminin α3 in nTERT keratinocytes by siRNA

Figure S2. Knockdown of Laminin332 in nTERT keratinocytes by lentiviral shRNA

Figure S3. Cholesterol biosynthesis genes in Collagen VII and Laminin-α3 knockdown 3D equivalents.

Figure S4. Lipidomic Analysis of *Lama3*flox/flox/K14-CreERT mouse epidermis

Figure S5. Lipidomic analysis of Laminin332 knockdown skin equivalents

Figure S6. LDL binding and internalisation.

Figure S7. Plasma Membrane Cholesterol in shLAMB3 and shLAMC2

Figure S8. Integrin-β1 and Focal Adhesion changes with loss of Lam332

Table 1. siRNA SMARTpool Tagret sequences

Table 2. DAVID GO Terms

Table 3. Junctional Epidermolysis Bullosa Patient details

Table 4. shRNA Clone target sequences

Table 5. qPCR primers and their sequences

Table 6. Antibodies and experimental conditions

Table 7. Internal standard mixture SPLASH Lipidomix® components

Movie S1. LDL movement in shC Control

Movie S2. LDL movement in shLAMA3 Control

Movie S3. LDL movement in shLAMB3 Control

Movie S4. LDL movement in shLAMC2 Control

Movie S5. LDL movement in shC on hrLaminin-332

Movie S6. LDL movement in shLAMA3 on hrLaminin-332

Movie S7. LDL movement in shLAMB3 on hrLaminin-332

Movie S8. LDL movement in shLAMC2 on hrLaminin-332

Movie S9. EGF movement in shC Control

Movie S10. EGF movement in shLAMA3 Control

Movie S11. EGF movement in shLAMB3 Control

Movie S12. EGF movement in shLAMC2 Control

## Abbreviations

JEB: Junctional Epidermolysis Bullosa
EB: Epidermolysis Bullosa
RDEB: Recessive Dystrophic Epidermolysis Bullosa
Lam332: Laminin-332
siLamA3: siRNA transfected keratinocytes
shLAMA3: shRNA transduced keratinocytes with Laminin-α3
shLAMB3: shRNA transduced keratinocytes with Laminin-β3
shLAMC2: shRNA transduced keratinocytes with Laminin-γ2
LC-MS: liquid chromatography -mass spectrometry
GC-MS: Gas chromatography -mass spectrometry
CE: Cholesterol ester
CS: Cholesterol sulfate
TC: total cholesterol
LDL: low density lipoprotein cholesterol
hrLam332: human recombinant Laminin-332
PM: Plasma membrane

