## Supplementary Figures and Tables for "Loss of cholesterol in Junctional Epidermolysis Bullosa skin identifies a key role for Laminin-332 in actomyosin mediated cholesterol transport"

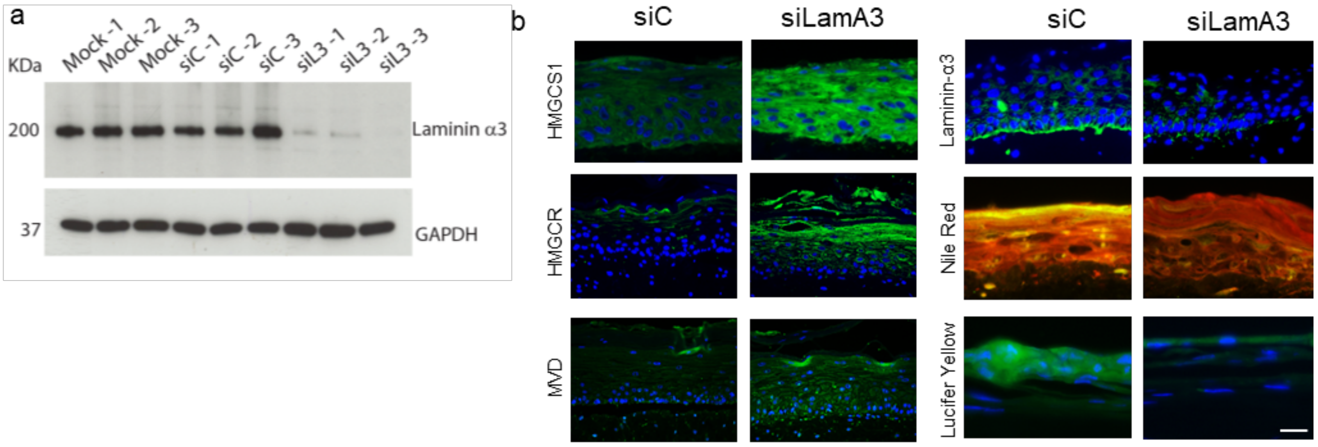


**Supplementary Figure 1. Knockdown of Laminin α3 in nTERT keratinocytes by siRNA**

**a)** Western blot analysis of Laminin α3 expression in nTERT keratinocytes transduced with small interfering RNA (siRNA) mock, siControl and siLaminin-α3, GAPDH was used as an internal control for protein loading. **b)** Immunofluorescence staining of cholesterol biosynthesis genes were performed in 3D skin equivalents and **c)** further analysis of barrier lipids with nile red stain (polar lipids in red and non-polar in green) and barrier function using lucifer yellow dye.

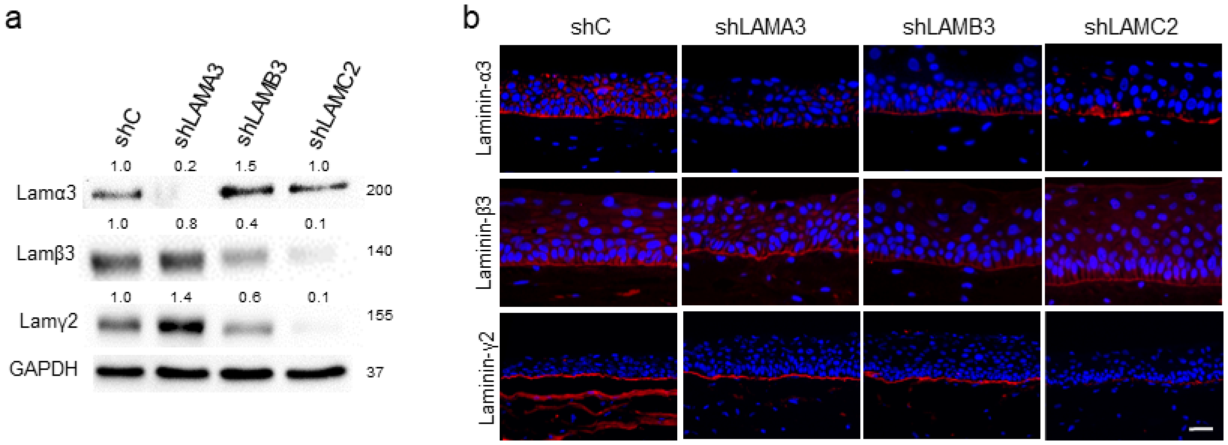


**Supplementary Figure 2. Knockdown of Laminin332 in nTERT keratinocytes by lentiviral shRNA**

**a)** Western blotting analysis of Lam332 expression in nTERT keratinocytes stably transduced with SMARTvector™ lentiviral particles of short hairpin ribonucleic acid (shRNA) clones (shLAMA3, shLAMB3 and shLAMC2) targeting each individual chain of Lam332. Non-targeting shRNA particles (shC) were used as a negative control. GAPDH was used as an internal control for protein loading. For densitometric analysis, results were normalized to GAPDH and are expressed as fold induction over shC. **b)** Immunofluorescence staining of sections of shC, shLAMA3, shLAMB3 and shLAMC2 3D skin equivalents showing loss of each individual Lam332 chain at the basement membrane. Scale bar = 50μm

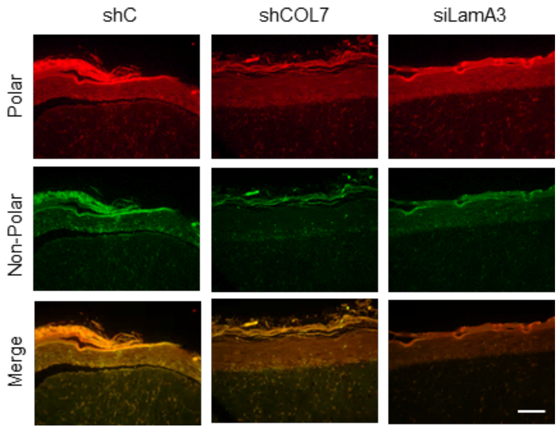


**Supplementary Figure 3. Nile Red Lipid staining in Collagen VII and Laminin-α3 knockdown 3D equivalents.**

Analysis of barrier lipids with nile red stain (polar lipids in red and non-polar in green) in shControl, shCol7 and siLamA3 in 3D skin equivalents.

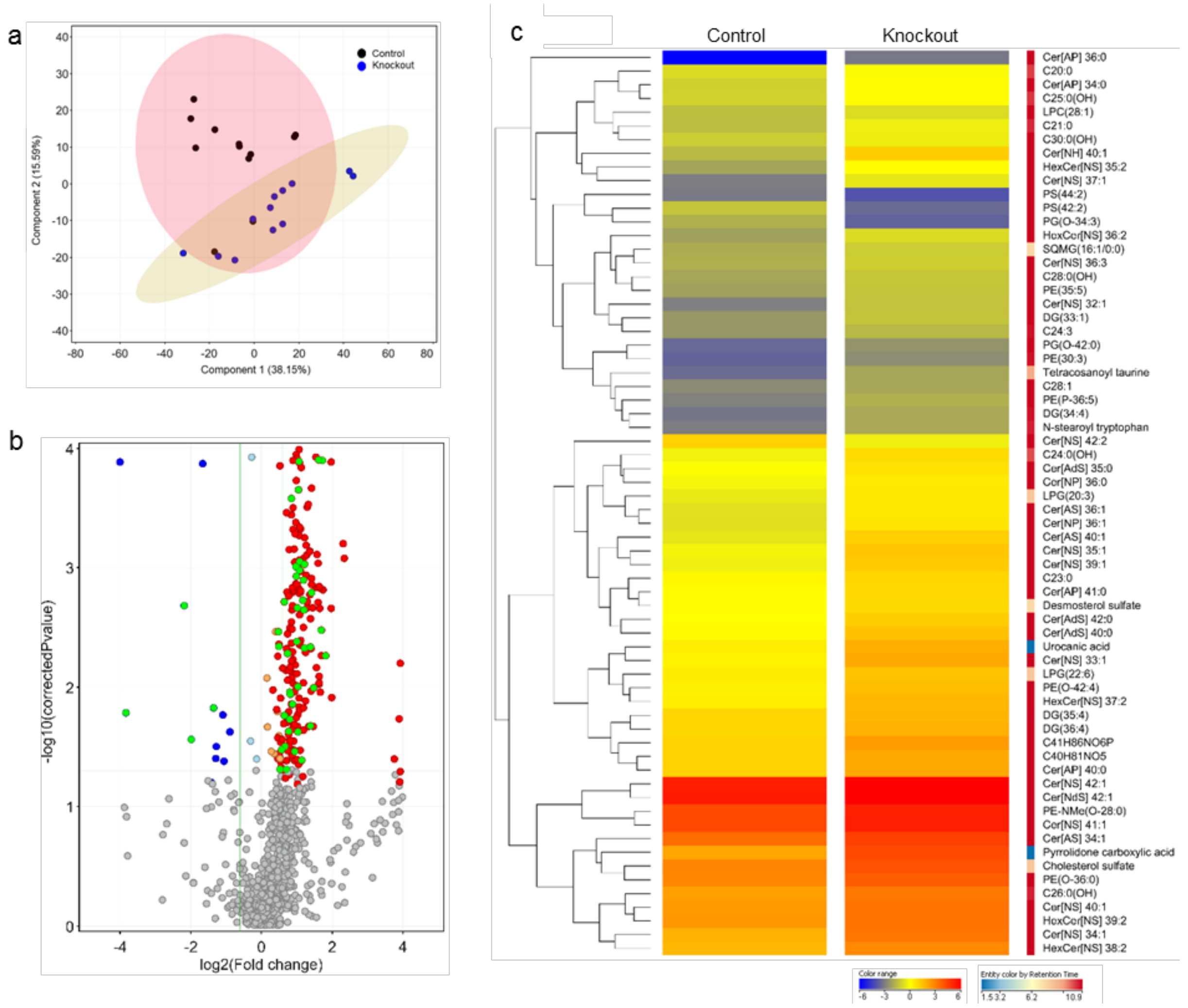


**Supplementary Figure 4. Lipidomic Analysis of *Lama3*$^{flox/flox}$/K14-CreERT Mouse epidermis.**

Analysis in Negative Ion mode (-ESI). **a)** Principal component analysis (PCA) illustrating the variances between the two sample groups. Within the first two principal components 53.74% of the total variance was explained, with 38.15% in the first dimension and an additional 15.59% in the second dimension. **b)** Volcano plot analysis displaying significantly different regulated lipid species between Lama3 KO vs Ctrl. The log2 of the FC values were plotted on the x-axis, whereas the −log10 of the t-test p-values were plotted on the y-axis. The dots corresponding to each entity were coloured according to the FC, with red indicating significantly upregulated lipids and blue indicating significantly downregulated (based on $p < 0.05$ and FC > 1.5); green dots highlight the lipid molecules further analysed in figure c. **c)** Hierarchical clustering analysis of 66 lipid species through targeted MS/MS which were significantly different with a FC >1.5 between KO and Ctrl. Colour scale (blue to red) reflects Log 2 values and colour scale (blue to red) shows retention time in the LC-MS analysis indicated in minutes.

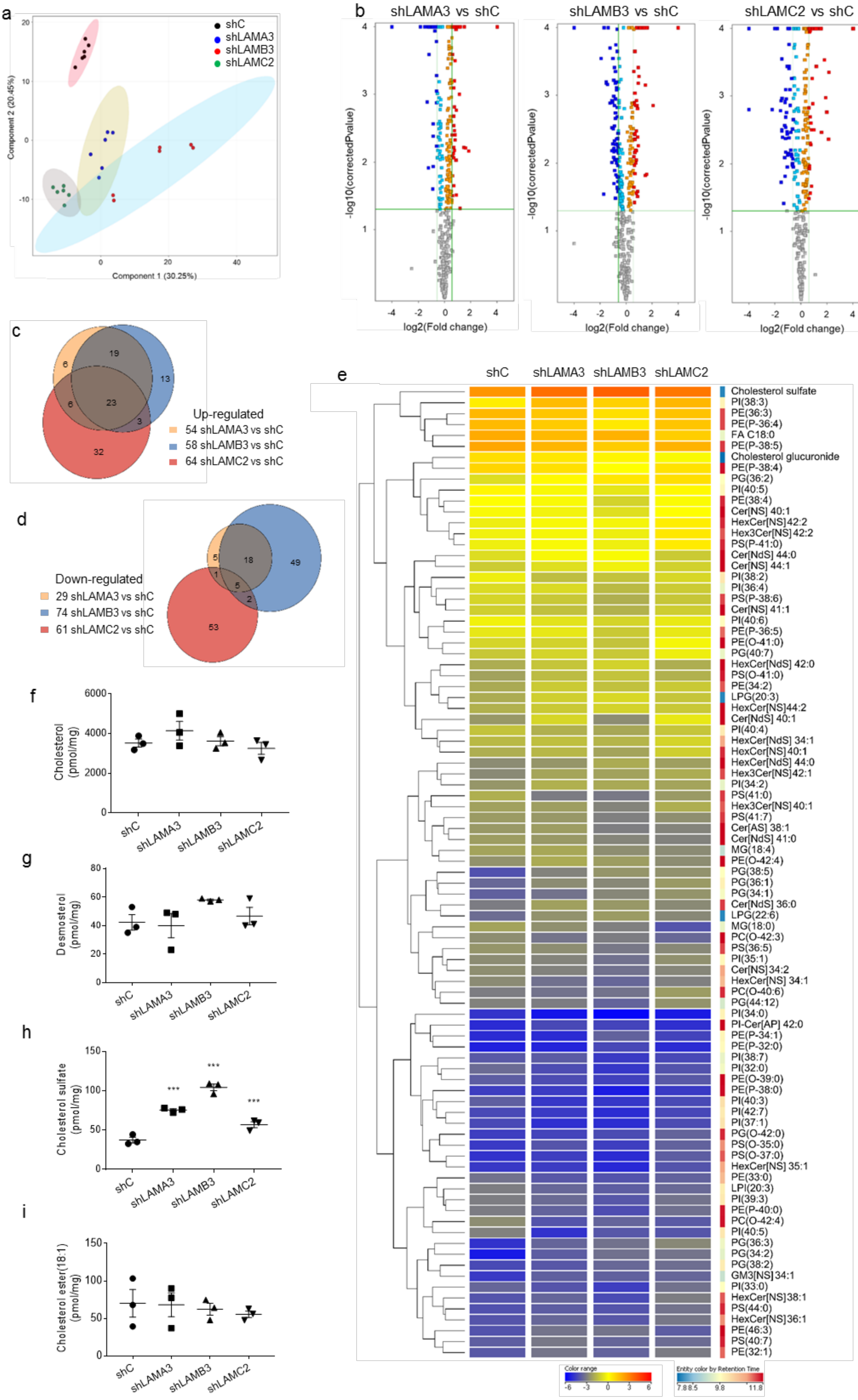
a
shC
shLAMA3
shLAMB3
shLAMC2
Component 2 (20.45%)
Component 1 (30.25%)
b
shLAMA3 vs shC
shLAMB3 vs shC
shLAMC2 vs shC
-log10(correctedPvalue)
log2(Fold change)
c
19
6
13
6
23
3
32
Up-regulated
54 shLAMA3 vs shC
58 shLAMB3 vs shC
64 shLAMC2 vs shC
d
Down-regulated
29 shLAMA3 vs shC
74 shLAMB3 vs shC
61 shLAMC2 vs shC
5
18
49
1
5
2
53
e
shC
shLAMA3
shLAMB3
shLAMC2
Cholesterol sulfate
PI(38:3)
PE(36:3)
PE(P-36:4)
FA C18:0
PE(P-38:5)
Cholesterol glucuronide
PE(P-38:4)
PG(36:2)
PI(40:5)
PE(38:4)
Cer[NS] 40:1
HexCer[NS]42:2
Hex3Cer[NS]42:2
PS(P-41:0)
Cer[NdS] 44:0
Cer[NS] 44:1
PI(38:2)
PI(36:4)
PS(P-38:6)
Cer[NS] 41:1
PI(40:6)
PE(P-36:5)
PE(O-41:0)
PG(40:7)
HexCer[NdS] 42:0
PS(O-41:0)
PE(34:2)
LPG(20:3)
HexCer[NS]44:2
Cer[NdS] 40:1
PI(40:4)
HexCer[NdS] 34:1
HexCer[NS]40:1
HexCer[NdS] 44:0
Hex3Cer[NS]42:1
PI(34:2)
PS(41:0)
Hex3Cer[NS]40:1
PS(41:7)
Cer[AS] 38:1
Cer[NdS] 41:0
MG(18:4)
PE(O-42:4)
PG(38:5)
PG(36:1)
PG(34:1)
Cer[NdS] 36:0
LPG(22:6)
MG(18:0)
PC(O-42:3)
PS(36:5)
PI(35:1)
Cer[NS]34:2
HexCer[NS] 34:1
PC(O-40:6)
PG(44:12)
PI(34:0)
PI-Cer[AP] 42:0
PE(P-34:1)
PE(P-32:0)
PI(38:7)
PI(32:0)
PE(O-39:0)
PE(P-38:0)
PI(40:3)
PI(42:7)
PI(37:1)
PG(O-42:0)
PS(O-35:0)
PS(O-37:0)
HexCer[NS] 35:1
PE(33:0)
LPI(20:3)
PI(39:3)
PE(P-40:0)
PC(O-42:4)
PI(40:5)
PG(36:3)
PG(34:2)
PG(38:2)
GM3[NS]34:1
PI(33:0)
HexCer[NS]38:1
PS(44:0)
HexCer[NS]36:1
PE(46:3)
PS(40:7)
PE(32:1)
Color range
Entity color by Retention Time
f
Cholesterol (pmol/mg)
g
Desmosterol (pmol/mg)
h
Cholesterol sulfate (pmol/mg)
***
i
Cholesterol ester(18:1) (pmol/mg)
shC
shLAMA3
shLAMB3
shLAMC2

**Supplementary Figure 5. Lipidomic analysis of Laminin332 knockdown skin equivalents.**

**a)** Principal component analysis (PCA) illustrating the variances between the four sample groups (shC, shLAMA3, shLAMB3 and shLAMC2). Within the first two principal components 50.70% of the total variance was explained, with 30.25% in the first dimension and an additional 20.45% in the second dimension. **b)** Volcano plots of entities found differently regulated in Lam332 skin equivalents. Comparison of relative abundance of lipid entities between shLAMA3 and shC, shLAMB3 and shC, and shLAMC2 and shC were performed by applying univariate significance analysis to #entities found in 100% of samples belonging to at least one condition. The log2 of the FC values were plotted on the x-axis, whereas the −log10 of the t-test p-values were plotted on the y-axis. The dots corresponding to each entity were coloured according to the FC, with red indicating significantly upregulated lipids and dark blue indicating significantly downregulated (based on $p < 0.05$ and FC > 1.5). Grey, yellow and light blue dots represent lipid entities that did not reach significance. Venn's diagram depicting the number of entities that were commonly modified when comparing each Lam332 knockdown to control **c)** shows upregulated lipid entities and **d)** shows down-regulated lipid entities. **e)** Hierarchical clustering of the 89 lipid entities that were commonly modified in each Lam332 knockdown compared to control through targeted MS/MS which were significantly different with a FC >1.5. Colour scale (blue to red) reflects Log 2 values colour scale (blue to red) shows retention time in the LC-MS analysis indicated in minutes. **f-i)** Quantitative analysis of cholesterol related lipid species, cholesterol, desmosterol (with GC-MS), cholesterol sulfate and cholesterol ester (CE(18:0)) (with LC-MS) respectively. Statistical analysis was performed using an unpaired t-test **$p < 0.01$ (n=3).

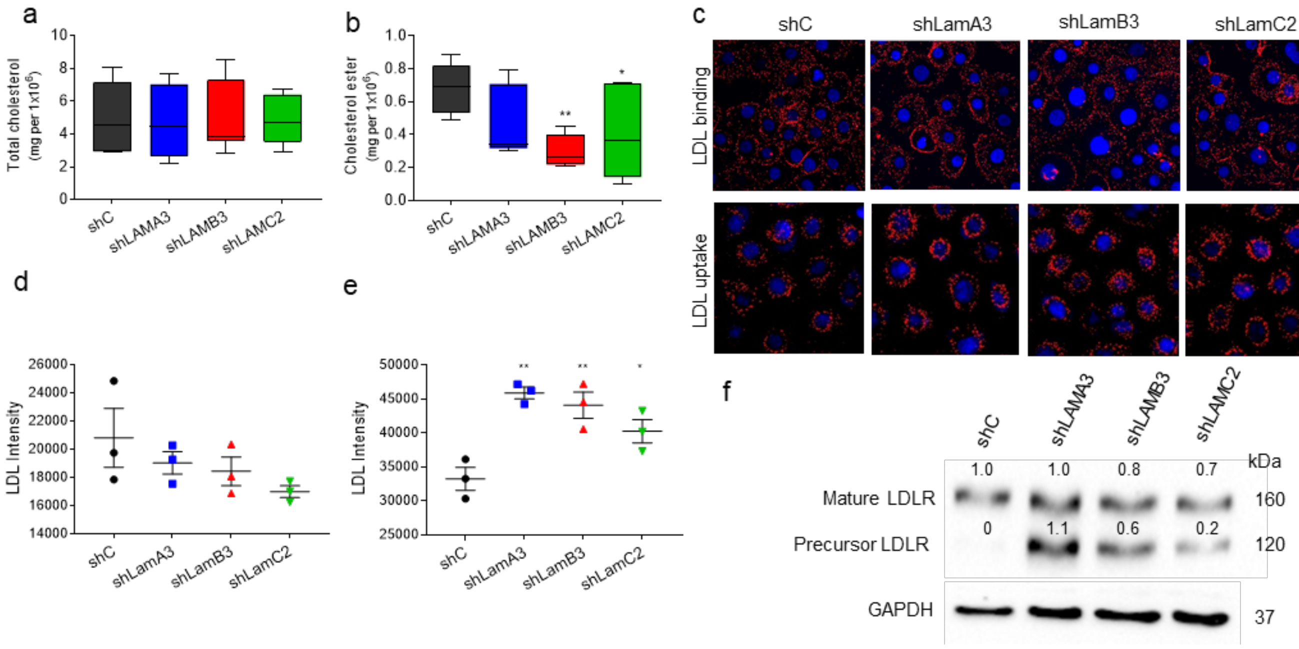


**Supplementary Figure 6. LDL binding and internalisation.**

**a)** Binding and cellular uptake of dil-LDL in control and Lam332 knockdown cells. Post-fixation images were taken using Incell 2200 and LDL fluorescence intensity per cell quantified using software, results shown for **b)** LDL binding and **c)** LDL internalisation respectively. Western blot analysis of LDL-receptor in Lam332 knockdown cells, showing both mature and precursor form of the receptor. GAPDH was used as an internal control for protein loading. For densitometric analysis, results were normalized to GAPDH and are expressed as fold induction over shC.

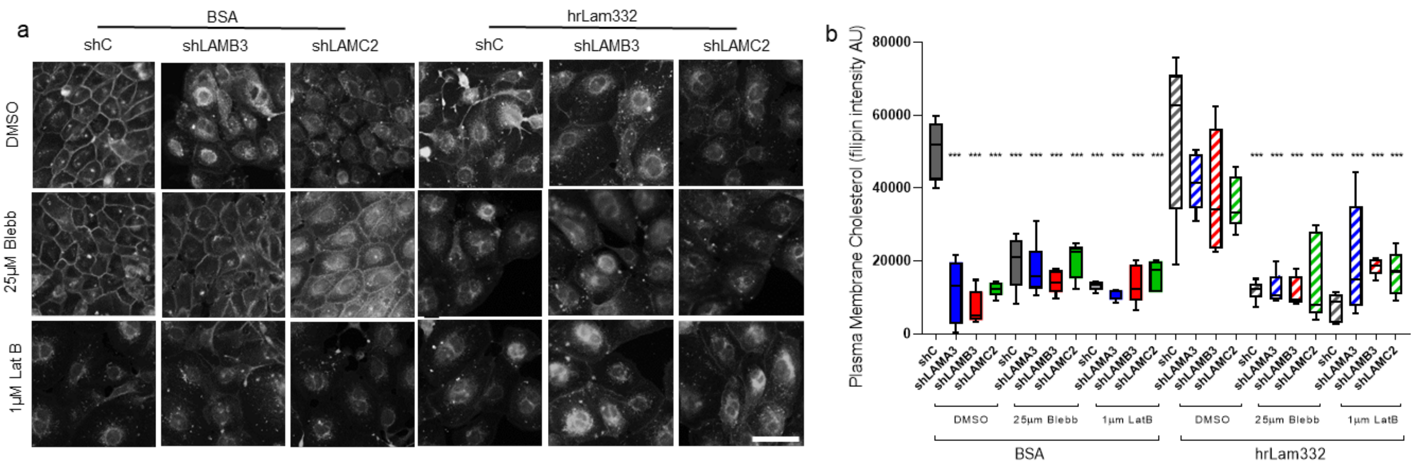


**Supplementary Figure 7. Plasma Membrane Cholesterol in shLAMB3 and shLAMC2**

**a)** Fluorescent filipin staining in control and shLAMB3 and shLAMC2 knockdown keratinocytes cultured on BSA or hrLam332, with and without Blebbistatin and Latrunculin B. **b)** Images were quantified and results presented as percentage of plasma membrane cholesterol. Graph shows analysis of all Lam332 knockdown cells (shLAMA3, shLAMB3 and shLAMC2) compared to control (shC). Statistical analysis was performed using one-way ANOVA, ***p<0.001. Scale bar 50µm.

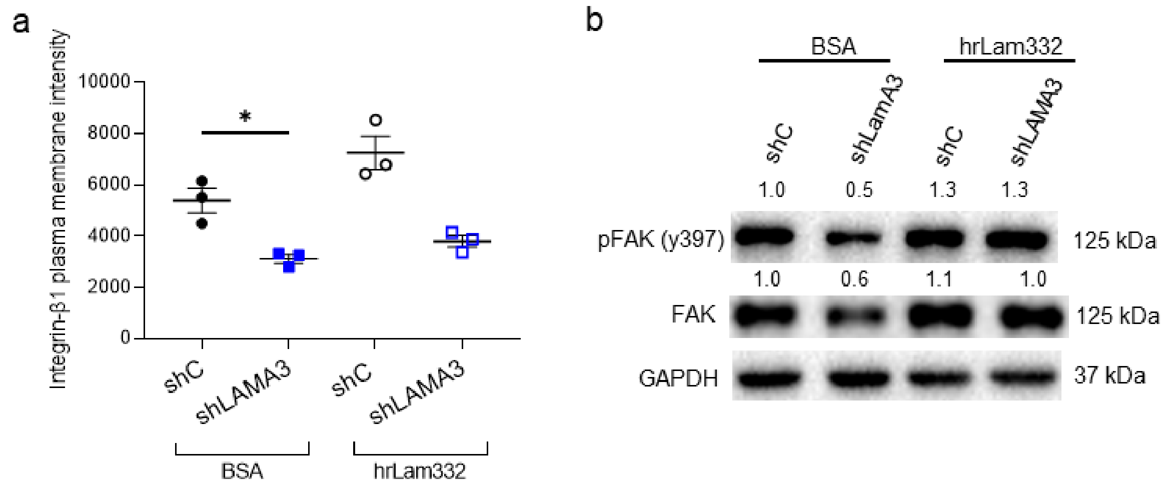


**Supplementary Figure 8. Integrin-β1 and Focal Adhesion changes with loss of Lam332**

**a)** Quantification of cellular plasma membrane Integrin-β1 in shLAMA3 and shC with cells cultured on BSA or hrLam332. Following flow cytometry analysis, the geometric mean of plasma membrane integrin-β1 intensity of single cells was calculated using the FlowJo software (v. 8.0) and corrected for the non-specific background signal using the isotype controls and DAPI only samples as negative controls. **b)** Western blot analysis of pFAK (y397) and total FAK in shLamA3 and shC with cells cultured on BSA or hrLam332 with GAPDH used as a loading control. Statistical analysis was performed using one-way ANOVA with Dunnett's multiple comparisons test with *p<0.05.

### Supplementary Tables

#### Supplementary Table 1. siRNA SMARTpool Target sequences.

| siRNA | Target Sequence | Reference |
|---|---|---|
| **siControl** | | |
| **siLAMA3** | GAAUUGAGCACCAGCGAUA<br>CAAUUGAGUUUCACUGAUU<br>GUCGUUAGAUUGAAUGAUA<br>GCAUAUGUGUUAACUGUCA | SO-3061252G |

#### Supplementary Table 2. DAVID GO Terms

| GO Term | PValue | Fold Enrichment | FDR |
|---|---|---|---|
| GO:0006695~cholesterol biosynthetic process | 1.03E-08 | 27.38462 | 1.68E-05 |
| GO:0008203~cholesterol metabolic process | 7.34E-08 | 10.6413 | 1.19E-04 |
| GO:0016126~sterol biosynthetic process | 9.72E-08 | 20.34286 | 1.58E-04 |
| GO:0016125~sterol metabolic process | 1.79E-07 | 9.693069 | 2.92E-04 |
| GO:0008610~lipid biosynthetic process | 3.25E-06 | 4.408669 | 0.005292 |
| GO:0006694~steroid biosynthetic process | 4.66E-06 | 9.423529 | 0.007578 |
| GO:0008202~steroid metabolic process | 1.63E-05 | 5.287129 | 0.026475 |
| GO:0008299~isoprenoid biosynthetic process | 0.001352 | 17.8 | 2.177409 |
| GO:0006720~isoprenoid metabolic process | 0.012925 | 8.090909 | 19.0789 |

#### Supplementary Table 3. Junctional Epidermolysis Bullosa Patient details

| Patient ID | Gene | Mutation |
|---|---|---|
| JEB Skin (*LAMA3)*-1 | LAMA3 | LAMA3. +/- c.6808 C>G, p.R2270X, exon 54 |
| JEB Skin (*LAMA3)*-2 | LAMA3 | |
| JEB Skin (*LAMA3)*-3 | LAMA3 | LAMA3. +/- c.6808 C>G, p.R2270X, exon 54 |
| JEB Skin (*LAMA3)-4* | LAMA3 | LAMA3. (+/+) c.8407ins4(ACCC), p.S2803fsX24, X65 PflMI/ DraIII cut mutnt sequence |
| JEB Skin (*LAMB3)*-1 | LAMB3 | LAMB3. c.1903 C>T, p.R635X, exon 14, c.978delC; p.H326fsX10 |
| JEB Skin (*LAMB3)*-2 | LAMB3 | LAMB3. (+/-) c.727 C>T; p.Q243X, X8 // (+/-) c.1903 C>T, pR635X, X14 |
| JEB Skin (*LAMB3)*-3 | LAMB3 | LAMB3. (+/) IVS8+1G>C AND (+/-) p.R635X |

**Supplementary Table 4. shRNA Clone target sequences**

| Target gene | Target Sequence | Catalogue # | LOT# |
|---|---|---|---|
| LAMA3 | GACGTATATGGATGGTTTA | SH-011071-03 | LV230912 |
| LAMB3 | GAGGCTACTGTAATCGCTA | SH-011072-02-10 | BV041201 |
| LAMC2 | GTGTATCTTTGATCGGGAA | SH-012119-01-10 | BV041201 |
| Non-Targeting negative | | S-005000-01 | GV251103 |

**Supplementary Table 5. qPCR primers and their sequences**

| Gene symbol | Forward primer (Sequence 5'-3') | Reverse primer (Sequence 5'-3') |
|---|---|---|
| *DHCR24* | CGTGTTGCCTGAGCTTGATG | AGTTTTCGGACGGAGTGCAT |
| *DHCR7* | AGGTGTGCGCAGGACTTTAG | CTTCTTGAACCGGCCCCTTA |
| *FDPS* | GAGACCGGGCCTTACTTCTG | GGACAGGGGCATCCTTCAAA |
| *HMGCR* | CTAGTGAGATCTGGAGGATCCAA | ACAAAGAGGCCATGCATTCG |
| *HMGCS1* | GGTGGGTTGGCGGCTATAAA | CTTCGGGCACAAGCGTGA |
| *LSS* | GTACGAGCCCGGAACATTCT | CCAGTCAGGAAACAGCCACA |
| *MVD* | TCAAGTACTGGGGCAAGCG | CAAATCCGGTCCTCGGTGAA |
| *NSDHL* | CGCCTACGGACGGAAAAGA | CGTGCGACTTGGTCTCTCAT |
| *HPRT* | GAAGAGCTATTGTAATGACC | GCGACCTTGACCATCTTTG |

**Supplementary Table 6. Antibodies and experimental conditions**

| Antigen | Host | Company | Dilution | | | |
|---|---|---|---|---|---|---|
| | | | WB | IF | ICC | FC |
| **phosphoFAK** | Mouse | BD Transduction Laboratories<br>611722 | 1:1000 | | | |
| **FAK** | Rabbit | Cell Signalling 3285 | 1:1000 | | | |
| **GAPDH** | Rabbit | Abcam ab9485 | 1:5000 | | | |
| **HMGCR** | Rabbit | Abcam ab174830 | 1:1000 | | | |

| | | | | | | |
|---|---|---|---|---|---|---|
| **HMGCR** | Rabbit | Abcam ab214018 | | 1:100 | 1:100 | |
| **HMGCS1** | Rabbit | Abcam ab155787 | 1:1000 | 1:100 | 1:100 | |
| **Integrin-β1** | Mouse | BD Pharminogen 559883 | | | | 1:100 |
| **Laminin-α3** | Mouse | Sigma | 1:1000 | | | |
| **Laminin-α3** | Rabbit | Atlas antibodies HPA009309 | | 1:100 | | |
| **Laminin-β3** | Mouse | SantaCruz 135968 | 1:1000 | 1:100 | | |
| **Laminin-γ2** | Mouse | Millipore mab19562 | 1:1000 | | | |
| **Laminin-γ2** | Rabbit | Proteintech 19698-1-ap | | 1:100 | | |
| **LSS** | Rabbit | gifted | 1:500* | 1:50 | 1:50 | |
| **MVD** | Rabbit | Novusbio NBP1-33050 | 1:500 | 1:50 | 1:50 | |
| **Myo5b** | Rabbit | Atlas antibodies (HPA040593) | 1:1000 | | 1:200 | |
| **NPC1** | Rabbit | Abcam 106534 | 1:1000 | | 1:200 | |
| **NPC2** | Rabbit | Abcam 218192 | 1:2000 | | 1:200 | |
| **Phalloidin** | | Invitrogen A22287 | | | 1:500 | |
| **Rab8a** | Rabbit | Abcam ab188574 | 1:6000 | | 1:1000 | |
| **Rab11a** | Rabbit | Invitrogen 71-5300 | 1:1000 | | 1:500 | |
| **α-Tubulin** | Mouse | Abcam ab7291 | 1:6000 | | 1:2000 | |

* Antibodies required 48H incubation

**Supplementary Table 7. Internal standard mixture SPLASH Lipidomix® components**

| Supplier | Country | SPLASH® Lipidomix® | Label | Chemical Formula | µmol/L | µL added to samples | pmole added to the sample |
|---|---|---|---|---|---|---|---|
| Avanti Polar Lipids | USA | 15:0-18:1(d7) PC | d7PC (33:1) | C41H73D7NO8P | 8.00 | 50 | 400 |
| Avanti Polar Lipids | USA | 15:0-18:1(d7) PE | d7PE (33:1) | C38H67D7NO8P | 0.30 | 50 | 15 |
| Avanti Polar Lipids | USA | 15:0-18:1(d7) PS (Na salt) | d7PS (33:1) | C39H66D7NNaO10P | 0.20 | 50 | 10 |
| Avanti Polar Lipids | USA | 15:0-18:1(d7) PG (Na salt) | d7PG (33:1) | C39H67D7NaO10P | 1.40 | 50 | 70 |
| Avanti Polar Lipids | USA | 15:0-18:1(d7) PI (NH4 salt) | d7PI (33:1) | C42H75D7NO13P | 0.40 | 50 | 20 |
| Avanti Polar Lipids | USA | 15:0-18:1(d7) PA | d7PA (33:1) | C36H61D7NaO8P | 0.40 | 50 | 20 |
| Avanti Polar Lipids | USA | 18:1(d7) LPC | d7LPC (18:1) | C26H45D7NO7P | 1.80 | 50 | 90 |
| Avanti Polar Lipids | USA | 18:1(d7) LPE | d7LPE (18:1) | C23H39D7NO7P | 0.40 | 50 | 20 |
| Avanti Polar Lipids | USA | 18:1(d7) Chol Ester | d7CE (18:1) | C45H71D7O2 | 20.0 | 50 | 1000 |
| Avanti Polar Lipids | USA | 18:1(d7) MG | d7MG (18:1) | C21H33D7O4 | 0.20 | 50 | 10 |
| Avanti Polar Lipids | USA | 15:0-18:1(d7) DG | d7DG (33:1) | C36H61D7O5 | 0.60 | 50 | 30 |
| Avanti Polar Lipids | USA | 15:0-18:1(d7)-15:0 TG | d7TG (48:1) | C51H89D7O6 | 2.60 | 50 | 130 |
| Avanti Polar Lipids | USA | 18:1(d9) SM | d9SM (d18:1/18:1) | C41H72D9N2O6P | 1.60 | 50 | 80 |
| Avanti Polar Lipids | USA | Cholesterol (d7) | d7Cholesterol | C27H39D7O | 10.0 | 50 | 500 |
| **Supplier** | **Country** | **Sphingolipid Mix I (LM 6002)** | **Label** | **Chemical Formula** | **µmol/L** | **µL added to samples** | **pmole added to the sample** |
| Avanti Polar Lipids | USA | Sphingosine (C17 base) | C17S | C17H35NO2 | 2.00 | 50 | 100 |
| Avanti Polar Lipids | USA | Sphinganine (C17 base) | C17DS | C17H37NO2 | 2.00 | 50 | 100 |
| Avanti Polar Lipids | USA | Sphingosine-1-P (C17 base) | C17S1P | C17H36NO5P | 2.00 | 50 | 100 |
| Avanti Polar Lipids | USA | Sphinganine-1-P (C17 base) | C17DS1P | C15H38NO5P | 2.00 | 50 | 100 |
| Avanti Polar Lipids | USA | 12:0 Sphingomyelin | SM(d18:1/12:0) | C35H71N2O6P | 2.00 | 50 | 100 |
| Avanti Polar Lipids | USA | 12:0 Ceramide | Cer(d18:1/12:0) | C30H59NO3 | 2.00 | 50 | 100 |
| Avanti Polar Lipids | USA | Glucosyl(ß) C12 Ceramide | HexCer(d18:1/12:0) | C36H69NO8 | 2.00 | 50 | 100 |
| Avanti Polar Lipids | USA | Lactosyl(ß) C12 Ceramide | Hex2Cer(d18:1/12:0) | C42H79NO13 | 2.00 | 50 | 100 |
| Avanti Polar Lipids | USA | 12:0 Ceramide-1-P | CerP(d18:1/12:0) | C30H60NO6P | 2.00 | 50 | 100 |
| Avanti Polar Lipids | USA | 25:0 Ceramide | Cer(d18:1/25:0) | C43H85NO3 | 2.00 | 50 | 100 |
| **Supplier** | **Country** | **Deuterated standards mixed in-house** | **Label** | **Chemical Formula** | **µmol/L** | **µL added to samples** | **pmole added to the sample** |
| Avanti Polar Lipids | USA | d31Cer(d18:1/16:0) | d31Cer[NS] | C34H36D31NO3 | 2.00 | 50 | 100 |
| C/D/N Isotopes | Canada | d17Palmitate | d17C16:0 | C16H15D17O2 | 100 | 50 | 5000 |
| C/D/N Isotopes | Canada | d4hexadecyl-d3hexadecanoate | d7WE (32:0) | C32H57D7O2 | 20.0 | 50 | 1000 |

| | | | | | | | |
|---|---|---|---|---|---|---|---|
| C/D/N Isotopes | Canada | Glyceryl trihexadecanoate-d98 | d98TG (48:0) | $C_{51}D_{98}O_6$ | 10.0 | 50 | 500 |
| C/D/N Isotopes | Canada | 5-Cholesten-3β-ol sulfate | d7CHS | $C_{27}H_{38}D_7NaO_4S$ | 20.0 | 50 | 1000 |
| Toronto Research Chemicals | Canada | d6Desmosterol | d6Desmosterol | $C_{27}H_{38}D_6O$ | 20.0 | 50 | 1000 |
